# Mapping of pseudouridine residues on cellular and viral transcripts using a novel antibody-based technique

**DOI:** 10.1101/2021.05.01.442255

**Authors:** Cecilia Martinez Campos, Kevin Tsai, David G. Courtney, Hal P. Bogerd, Christopher L. Holley, Bryan R. Cullen

## Abstract

Pseudouridine (Ψ) is the most common non-canonical ribonucleoside present on mammalian non-coding RNAs (ncRNAs), including rRNAs, tRNAs and snRNAs, where it contributes ∼7% of the total uridine level. However, Ψ constitutes only ∼0.1% of the uridines present on mRNAs and its effect on mRNA function remains unclear. Ψ residues have been shown to inhibit the detection of exogenous RNA transcripts by host innate immune factors, thus raising the possibility that viruses might have subverted the addition of Ψ residues to mRNAs by host pseudouridine synthase (PUS) enzymes as a way to inhibit antiviral responses in infected cells. Here, we describe and validate a novel antibody-based Ψ mapping technique called photo-crosslinking assisted Ψ sequencing (PA-Ψ-seq) and use it to map Ψ residues on not only multiple cellular RNAs but also on the mRNAs and genomic RNA encoded by HIV-1. We describe several 293T-derived cell lines in which human PUS enzymes previously reported to add Ψ residues to human mRNAs, specifically PUS1, PUS7 and TRUB1/PUS4, were inactivated by gene editing. Surprisingly, while this allowed us to assign several sites of Ψ addition on cellular mRNAs to each of these three PUS enzymes, the Ψ sites present on HIV-1 transcripts remained unaffected. Moreover, loss of PUS1, PUS7 or TRUB1 function did not significantly reduce the level of Ψ residues detected on total human mRNA below the ∼0.1% level seen in wild type cells, thus implying that the PUS enzyme(s) that adds the bulk of Ψ residues to human mRNAs remains to be defined.

## Introduction

Eukaryotic mRNAs are subject to a number of covalent modifications at the single nucIeotide level, collectively referred to as epitranscriptomic modifications, that can regulate the translation, stability, splicing and/or immunogenicity of the mRNA (Gilbert et al. 2016; Roundtree et al. 2017; Tsai and Cullen 2020). For example, methylation of the *N*^6^ position of adenosine gives rise to m^6^A, the most common epitranscriptomic modification of cellular mRNAs, representing ∼0.4% of all adenosines (Meyer and Jaffrey 2014; Yue et al. 2015). Interestingly, m^6^A is detected at even higher levels on viral mRNAs, where it has been proposed to enhance viral mRNA function resulting in increased virus replication (Courtney et al. 2019a; Courtney et al. 2019c; Tsai and Cullen 2020). Another common epitranscriptomic RNA modification, called pseudouridine and abbreviated as Ψ, is an isomer of uridine found on a wide range of non-coding RNA (ncRNA) classes, including tRNAs, snRNAs and rRNAs (Borchardt et al. 2020). While Ψ is the most common non-canonical base detected in eukaryotic cells, representing 7-9% of all uridine residues in total cellular RNA, Ψ is detected at more modest levels on cellular mRNAs, where it has been reported to represent ∼0.2-0.3% of all uridines (Hughes and Maden 1978; Li et al. 2015). However, Ψ is of considerable virological interest as Ψ has been reported to inhibit the detection of exogenous RNA molecules by host innate immune factors including Toll-like receptors (TLRs), retinoic acid-inducible gene I (RIG-I) and RNA-dependent protein kinase (PKR) (Kariko et al. 2005; Karikó et al. 2008; Anderson et al. 2010; Durbin et al. 2016). Overall, the presence of Ψ on exogenous mRNA molecules has been reported to not only prevent the induction of an interferon response but also increase mRNA stability and translation. As a result, synthetic mRNAs designed for use *in vivo* have Ψ, or more accurately N^1^-methylpseudouridine-exclusively in place of uridine, as seen for example in both the Moderna mRNA 1273 and the Pfizer/BioNTech BNT162b2 RNA vaccines for COVID-19 (Nance and Meier 2021). The reduced immunogenicity of mRNAs containing Ψ suggests that viruses might have co-opted cellular mechanisms that deposit Ψ on their mRNAs in order to avoid host innate immune responses and facilitate viral gene expression and replication. However, to our knowledge this question has not been previously addressed.

Ψ residues are deposited on RNAs via two distinct mechanisms. Ψ residues on rRNAs and some snRNAs, representing the majority of cellular Ψ, are deposited by small nucleolar ribonucleoproteins (snoRNPs) guided by base pairing between a box H/ACA snoRNA and the RNA target (Borchardt et al. 2020). Currently, no mRNA targets for Ψ addition via snoRNPs are known. The second mechanism for addition of Ψ to RNA transcripts depends on so-called stand-alone pseudouridine synthase (PUS) enzymes, of which 12 are known to exist in human cells (Rintala-Dempsey and Kothe 2017; Borchardt et al. 2020). Most PUS enzymes are localized either to mitochondria or nuclei and, in the latter case, are thought to modify RNAs co-transcriptionally (Martinez et al. 2020). Of the various PUS enzymes, PUS1 and PUS7 have been reported to be the predominant PUS enzymes for addition of Ψ to specific locations on mRNAs in yeast (Carlile et al. 2014; Schwartz et al. 2014) while in humans, Ψ has been reported to be added to mRNAs by not only PUS1 and PUS7 but also by TRUB1 (Li et al. 2015; Safra et al. 2017; Borchardt et al. 2020). However, at this point, few Ψ residues have been mapped on mRNAs expressed in human cells and for the majority of these, the responsible PUS enzyme remains undefined.

Here, we report the use of a novel antibody based Ψ mapping technology, called photo-crosslinking-assisted Ψ sequencing (PA-Ψ-seq), to map Ψ residues deposited on transcripts encoded by HIV-1 as well as on cellular mRNAs in HIV-1 infected cells. We report that HIV-1 transcripts are indeed modified by the addition of Ψ at specific locations and describe efforts to define the origin of these Ψ residues using human cell lines in which PUS1, PUS7 or TRUB1 had been knocked out by gene editing using CRISPR/Cas. Unexpectedly, we observed that none of the Ψ residues detected on HIV-1 transcripts were lost when any of these three human PUS enzymes were deleted and moreover loss of these enzymes did not significantly reduce the level of detection of Ψ residues on total human mRNA below the ∼0.1% level seen in wild type cells.

## Results

Previously, Ψ residues on cellular mRNAs have primarily been mapped using a chemical approach in which Ψ residues are selectively modified using N-cyclohexyl-N’(2-morpholinoethyl)-carbodiimide metho-p-toluenesulphonate (CMC), which induces a strong stop during reverse transcription (Carlile et al. 2015; Li et al. 2015; Safra et al. 2017; Zhang et al. 2019). While this technique has the power to potentially not only map Ψ residues with single nucleotide precision but also quantify the level of Ψ at any given site, it requires large amounts of input RNA and, even then, has the tendency to give rise to a significant level of both false positives and false negatives, as recently discussed (Wiener and Schwartz 2021). We therefore decided to develop a simpler, alternative technique based on antibody recognition of Ψ residues that, while not as precise as CMC-based techniques, nevertheless would allow the mapping of Ψ residues with high, ∼30nt precision using a much lower level of input mRNA.

We and others have described the use of photo-assisted crosslinking of specific residues on mRNAs to a cognate antibody, followed by deep sequencing (PA-mod-seq), to precisely map the locations of m^6^A, m^5^C and ac4C residues on viral and cellular RNAs (Chen et al. 2015; Kennedy et al. 2016; Courtney et al. 2019c; Tsai et al. 2020) and we therefore asked whether a commercial Ψ-specific antibody could be used to map Ψ residues. We first tested the accuracy of this novel technique, here referred to as PA-Ψ-seq, by using it to detect known Ψ residues on cellular ncRNAs and mRNAs. Human CEM T cells or 293T cells were pulsed overnight with a photoactivatable ribonucleoside, either 4-thiouridine (4SU) or 6-thioguanosine (6SG), and total RNA harvested (Hafner et al. 2010). Total RNA was then subjected to a single round of poly(A)+ RNA isolation using oligo-dT beads and the partially purified mRNA then mixed with an RNA modification-specific antibody and crosslinked using UV light. RNA-protein complexes were then isolated and incubated with RNAse T1. Protected RNA fragments were recovered by proteinase K treatment, ligated to adapters, reverse transcribed and subjected to deep sequencing. A valuable aspect of this technique is that crosslinked 4SU and 6SG both induce the specific misincorporation of residues during reverse transcription, G in the case of 4SU and T in the case of 6SG, thus allowing non-crosslinked reads to be selectively removed during bioinformatic analysis (Hafner et al. 2010). We initially used the PA-Ψ-seq technique to map known Ψ residues on cellular 28S rRNA (Fig. 1A), on the cellular mRNAs encoding RPL15 (Fig. 1B) or SRSF1 (Fig. 1C) as well as on tRNAs (represented by tRNA-Glu(CTC) in Fig. 1D) and U5 snRNA (Fig. 1E). Interestingly, this technology worked equally well using either 4SU or 6SG as the photoactivatable ribonucleoside, a result which is consistent with the finding that 4SU retains the ability to undergo isomerization by PUS enzymes (Zhou et al. 2010). Previously, Ψ residues on cellular mRNAs have been mapped predominantly to the coding sequence (CDS) as well as to the 3’ untranslated region (UTR) using the above referenced CMC mapping technique (Carlile et al. 2014; Li et al. 2015). In our hands, we found that the majority of Ψ residues actually mapped to the 3’UTR, including proximal to the translation termination codon (Supp. Fig. 1). While Ψ residues would not therefore be expected to have a major effect on codon usage, they could potentially affect the efficiency of translation termination, as has been previously suggested (Karijolich and Yu 2011).

**Figure 1.**
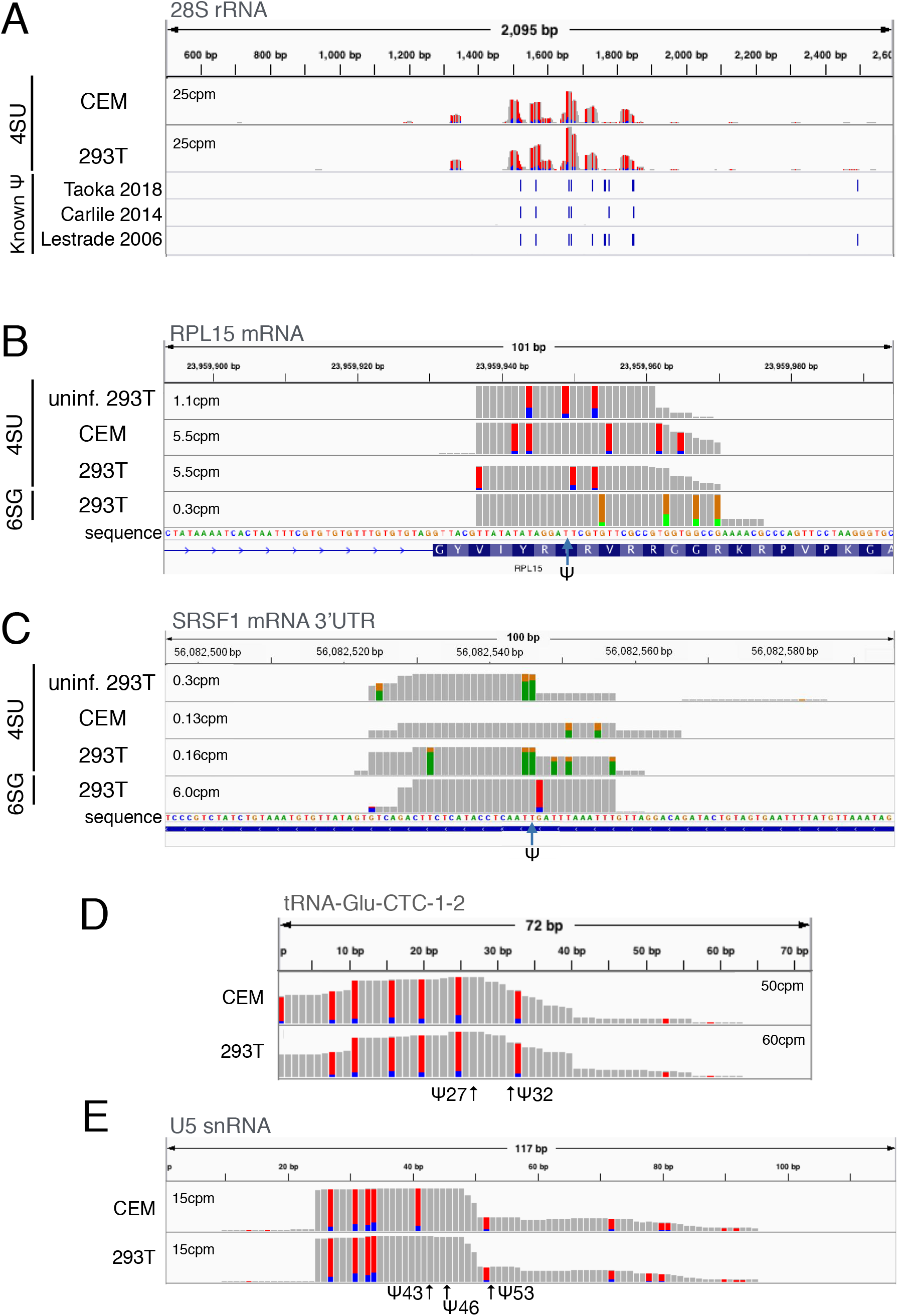
Validation of the novel PA-Ψ-seq mapping technique. The figure shows the location of previously published mapped Ψ sites on various ncRNAs and mRNAs. A) Mapping of Ψ residues on human 28S rRNA in CEM T cells and 293T cells pulsed with 4SU. Blue bars below the figure indicate the location of Ψ residues previously reported by Taoka *et al* 2018, Carlile *et al* 2014 and Lestrade and Weber, 2006, which show slight differences. B) Mapping of Ψ residues on human RPL15 mRNA in CEM T cells and 293T cells pulsed with 4SU or 6SG. The location of the single Ψ residue previously mapped in this mRNA is indicated by an arrow. C) Similar to panel B but mapping the single Ψ residue previously identified in SRSF1 mRNA. D) Mapping the two Ψ residues previously located in tRNA-Glu-CTC-1-2 in 4SU pulsed cells. E) Mapping the location of the three Ψ residues previously identified in U5 snRNA.

### Mapping Ψ residues on HIV-1 transcripts

We next asked if we could detect Ψ residues on the genomic RNA or mRNAs encoded by HIV-1. As shown in Fig. 2A, we were able to map at least 12 Ψ residues on viral transcripts in essentially every viral open reading frame (ORF) as well as in the 3’UTR. These Ψ residues were detected at the same locations in both virion and intracellular viral transcripts and on HIV-1 RNAs transcribed in CEM T cells and in 293T cells (Fig. 2A). Of note, none of the mapped Ψ residues coincided with known important viral RNA structures such as the apical region of TAR, the RRE or the Gag-Pol frameshift stem-loop.

**Figure 2.**
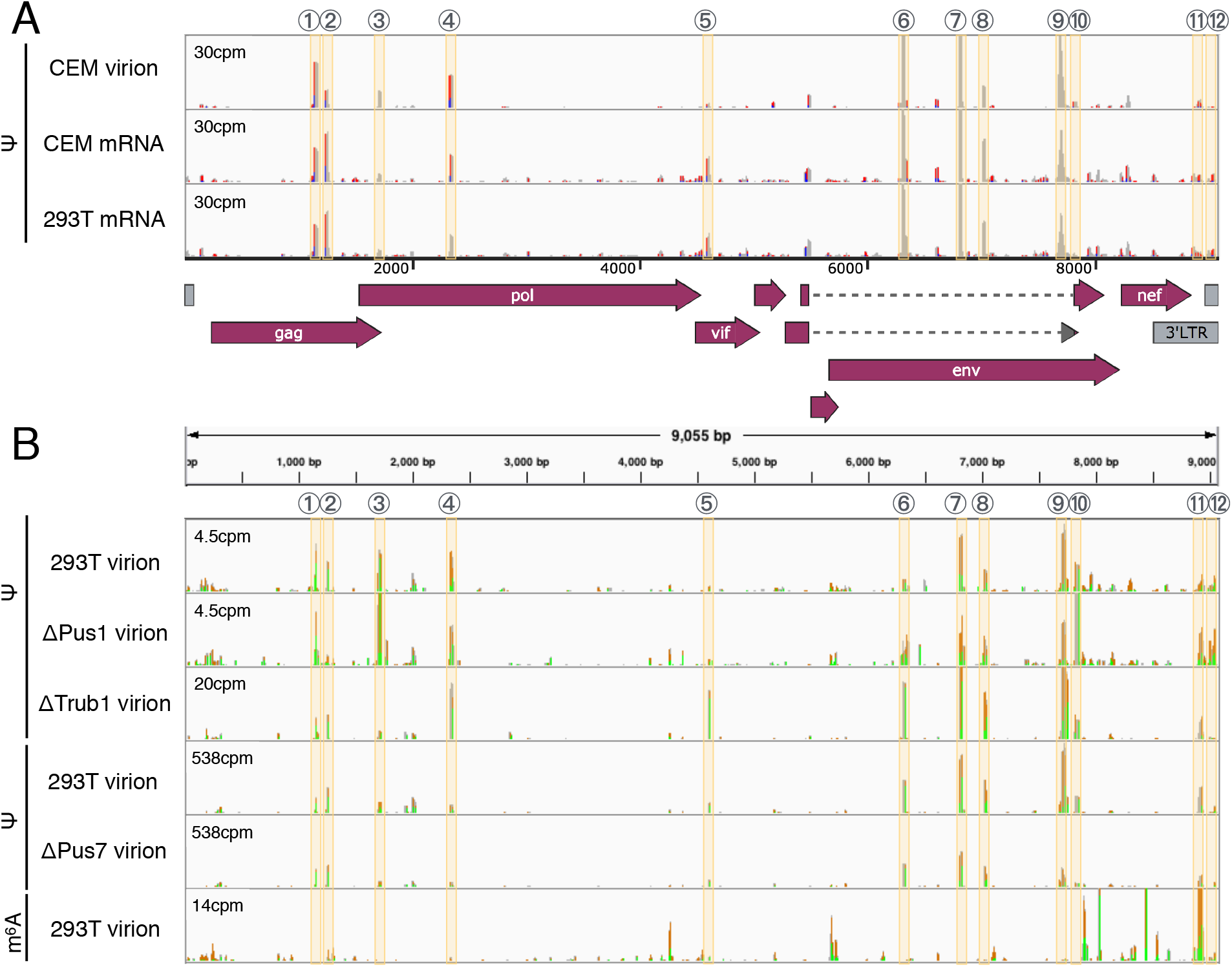
Using PA-Ψ-seq to map Ψ residues located on HIV-1 transcripts. A) HIV-1-infected CEM T cells or 293T cells were pulsed with 4SU and virion RNA produced by CEM cells, or poly(A)+ RNA isolated from CEM or 293T cells, analyzed for the presence of Ψ using PA-Ψ-seq B) Similar to panel A except that these data were obtained using total HIV-1 virion RNA isolated from virions released by 6SG-pulsed wild type or PUS-KO 293T cells, as indicated. All these data used the PA-Ψ-seq technique except for the last lane, which used an m6A-specific antibody to perform the very similar PA-m6A-seq technique as a specificity control. Highly reproducible Ψ peaks are numbered and indicated by beige lines and are shown relative to a schematic of the ORFs present on the HIV-1 genome.

### Derivation of 293T cell clones lacking individual PUS enzymes

The key question is of course whether Ψ is able to regulate retroviral gene expression and replication, as we hypothesize. However, because most mapped Ψ residues on HIV-1 transcripts are located in ORFs, only a subset could in principle be mutated without affecting the underlying amino acid sequence and, given the complex alternative splicing pattern that is characteristic of HIV-1, even apparently silent mutations might disrupt one of the many viral splicing enhancer or suppressor sequences and thereby inhibit viral gene expression by a Ψ-independent mechanism. We therefore decided to focus instead on identifying the cellular PUS enzyme(s) responsible for the modification of HIV-1 transcripts using a gene editing approach, as our analysis did not reveal any significant homology of the region flanking the mapped Ψ residues to known H/ACA snoRNAs (not shown). Previous work from others has identified PUS1, PUS7 and TRUB1 (known as PUS4 in yeast) as the main PUS enzymes responsible for adding Ψ residues to mammalian mRNAs (Li et al. 2015; Safra et al. 2017; Borchardt et al. 2020) and we therefore individually knocked out each of these enzymes by gene editing using CRISPR/Cas. The resultant polyclonal cell lines were then subjected to single cell cloning and two clonal cell lines in which PUS1, PUS7 or TRUB1 was knocked out identified by Western blot analysis (Fig. 3 A-C) combined with Sanger sequencing of the PCR amplified genomic DNA target site (Supp. Fig. 2). In none of the six clonal cell lines could the parental genomic sequence be recovered by PCR, thus confirming the absence of a functional PUS enzyme. Targeted gene editing resulted in the clonal cell line ΔPUS1-1, bearing a 22nt deletion in exon 2 as well as a 336nt deletion that removed all of exon 1, including the start codon, as well as ΔPUS1-2, bearing 5nt and 64nt deletions in exon 2. Similarly, the cell line ΔPUS7-1 had a 4nt and an 8nt deletion in exon 10 while ΔPUS7-2 had an 8nt deletion. Finally, ΔTRUB1-1 had a 220nt, a 219nt or a 231nt deletion in exon 1 while ΔTRUB1-2 had a 102nt and two 220nt deletions in exon 1, all of which removed part of the essential “motif 1” region in TRUB1 (Zucchini et al. 2003). Our ability to generate knock out cell lines for three different cellular PUS enzymes is perhaps surprising, as Ψ is thought to play a critical role in RNA metabolism and these PUS proteins potentially could have been essential. We did notice that all six clonal cell lines grew more slowly than the parental 293T cells and this effect was especially obvious for the two ΔPUS1 cell lines (Supp. Fig. 3).

**Figure 3.**
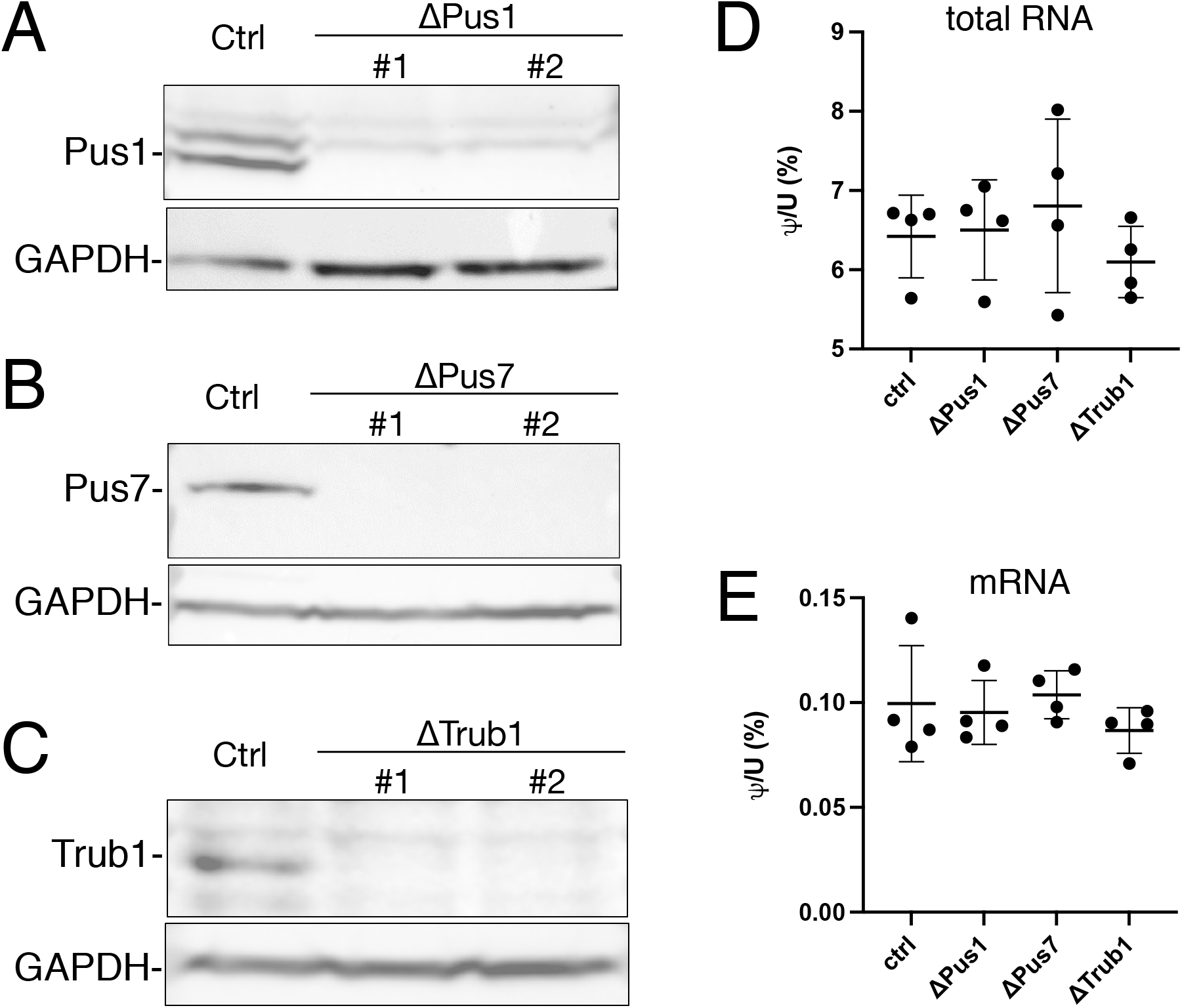
Validation of cellular PUS enzymes knock-out cell lines. A) Western blot of WT and two ΔPUS1 clonal cell lines B) Western blot of WT and two ΔPUS7 clonal cell lines C) Western blot of WT and two ΔTRUB1 clonal cell lines. D) Analysis of Ψ levels in total RNA samples derived from WT or PUS-deficient 293T clones using UPLC-MS/MS. N=4 biological replicates with each PUS knock out cell line analyzed twice. Individual values, average and SD indicated. E) Similar to panel D except analyzing highly purified mRNA samples derived from the indicated WT or PUS-deficient 293T clones.

### Quantification of Ψ levels in human mRNA

Armed with clonal cell lines lacking each of the cellular PUS enzymes PUS1, PUS7 or TRUB1, we asked whether any of these enzymes has an impact on the overall Ψ content of cellular RNA. To do this, we measured the Ψ content for both total cellular RNA and highly-purified mRNA, using ultra-high-performance liquid chromatography and tandem mass spectrometry (UPLC-MS/MS). Poly-adenylated mRNA was isolated by two rounds of oligo-dT affinity purification using oligo-dT, followed by a RiboZero to eliminate any residual rRNA, since rRNA is highly enriched for snoRNA-dependent Ψ modifications. mRNA purity was assessed by bioanalyzer and measurements of both m^6,6^A and t^6^A (which are thought to be present only on rRNA and tRNA respectively); both of these adenosine modifications were present at >100 fold the limit of detection in total RNA but undetectable in our mRNA samples (Supp. Fig. 4). As shown in Figure 3, none of the KO cell lines showed a significant loss of overall Ψ content in either total cellular RNA (Fig. 3D) or highly purified mRNA (Fig. 3E). Of note, the amount of Ψ content detected on highly purified mRNA from our cell lines (∼0.1%) was somewhat lower than previously reported by Li et al. (0.21%) (Li et al. 2015), but our method differs in that it incorporates two stable-isotope-labeled spike-in nucleoside standards, and our results may therefore represent a more accurate estimation of the true Ψ content. Moreover, we note that Li et al. reported a level of 0.0017% m^6,6^A in their purified mRNA samples, which is >4 fold higher that seen in our mRNA preparations (Supp. Fig. 4) and implies a level of contamination of ∼1% rRNA, which would be sufficient to increase the level of Ψ detected in their preparation by ∼0.07%

Although we did not detect any global changes to pseudouridine content in total cellular RNA or mRNA, we next asked if any of the mapped sites of Ψ addition on HIV-1 transcripts had been lost (Fig. 2B). However, we found that all 12 sites of Ψ addition on the HIV-1 RNA genome were still readily detectable in each of the KO cell lines. To further confirm the specificity of the PA-Ψ-seq technique, we performed the closely similar PA-m^6^A-seq technique using one of the same RNA samples (PA-m^6^A-seq is identical to PA-Ψ-seq except that an m^6^A-specific antibody is substituted for the Ψ-specific antibody) and observed the previously reported pattern of m^6^A sites on the HIV-1 RNA genome, with sites of m^6^A addition concentrated in the *nef* gene and the 3’ long terminal repeat (Kennedy et al. 2016). We next sought to identify sites of Ψ addition to cellular mRNAs that were lost in one of the KO 293T cell lines using a very stringent bioinformatic approach. In fact, we were able to identify 16 high confidence PUS1-dependent Ψ sites, 41 high confidence PUS7-dependent Ψ sites and 9 high confidence TRUB1 dependent sites on a range of cellular mRNAs (Supp. Table 1). Of these, the PA-Ψ–seq data for two of the PUS1-dependent sites are shown in Figs.4A and B, for two of the PUS7-dependent sites in Figs. 4C and D, and the data for two of the TRUB1-dependent sites in Figs. 4E and F.

**Figure 4.**
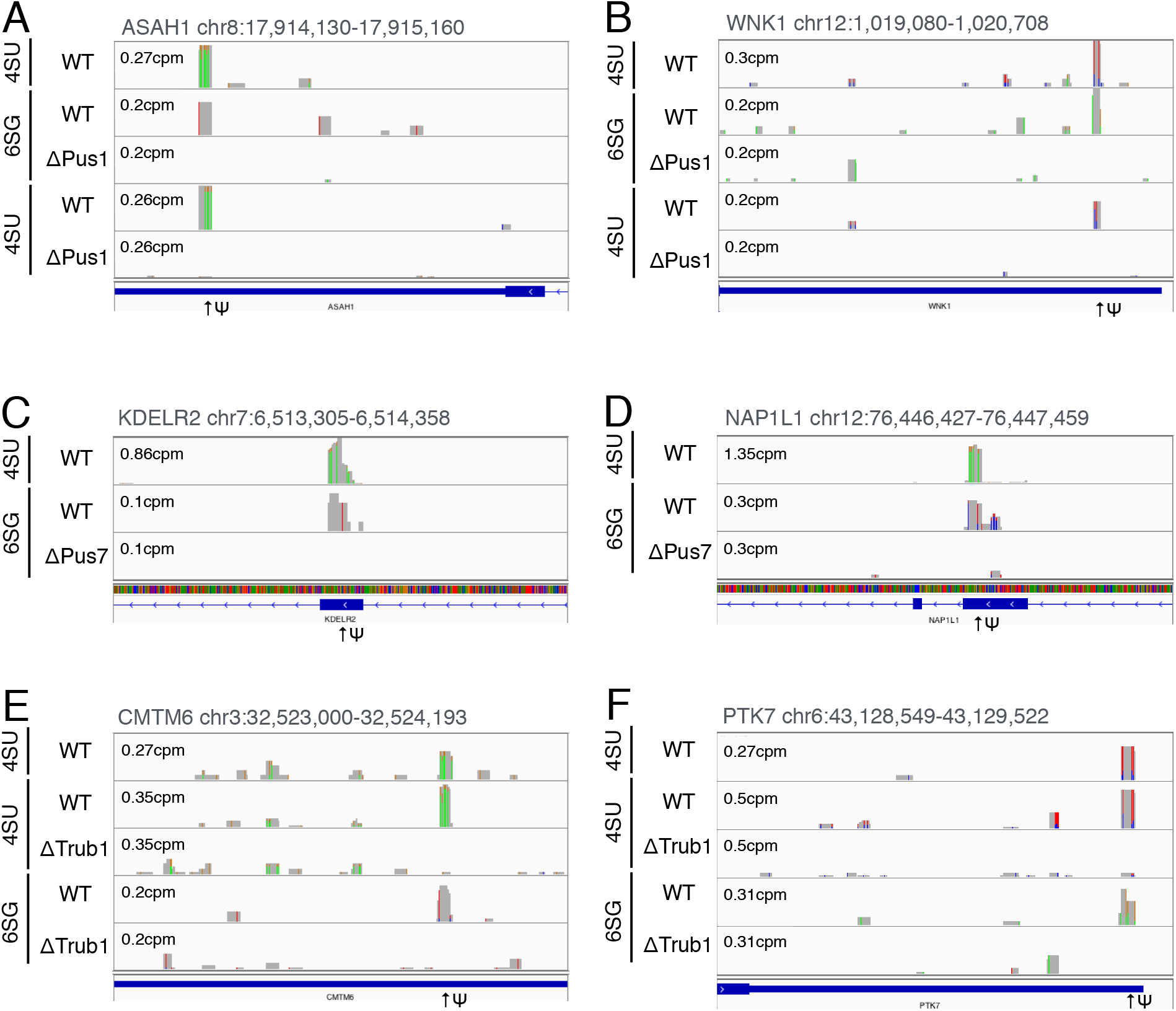
Representative PUS1, PUS7 and TRUB1-dependent Ψ residues on cellular mRNAs. Similar to figure 1 except that PA-Ψ-seq was used to map Ψ residues in poly(A)+ RNA isolated from WT 293T cells as well as from ΔPUS1 (panels A and B), ΔPUS7 (panels C and D) and ΔTRUB1 (panels E and F) clonal 293T cell lines. The location of Ψ residues in mRNA exonic locations (exons indicated by blue boxes) is indicated by arrows.

The unexpected finding that none of the mapped Ψ residues on the HIV-1 genome were lost in any of the three PUS KO cell lines meant that we could not use these cell lines to look at the effect in *cis* of Ψ residues on HIV-1 gene expression. Nevertheless, we reasoned that loss of one or more of these PUS enzymes might affect HIV-1 gene expression indirectly, for example by interfering with the expression of cellular RNAs. This hypothesis is tested in Figs. 5A and B, which show that loss of PUS1 expression does indeed result in a modest but highly significant ∼30% decline in HIV-1 Gag protein expression, while loss of either PUS7 or TRUB1 had no detectable effect.

**Figure 5.**
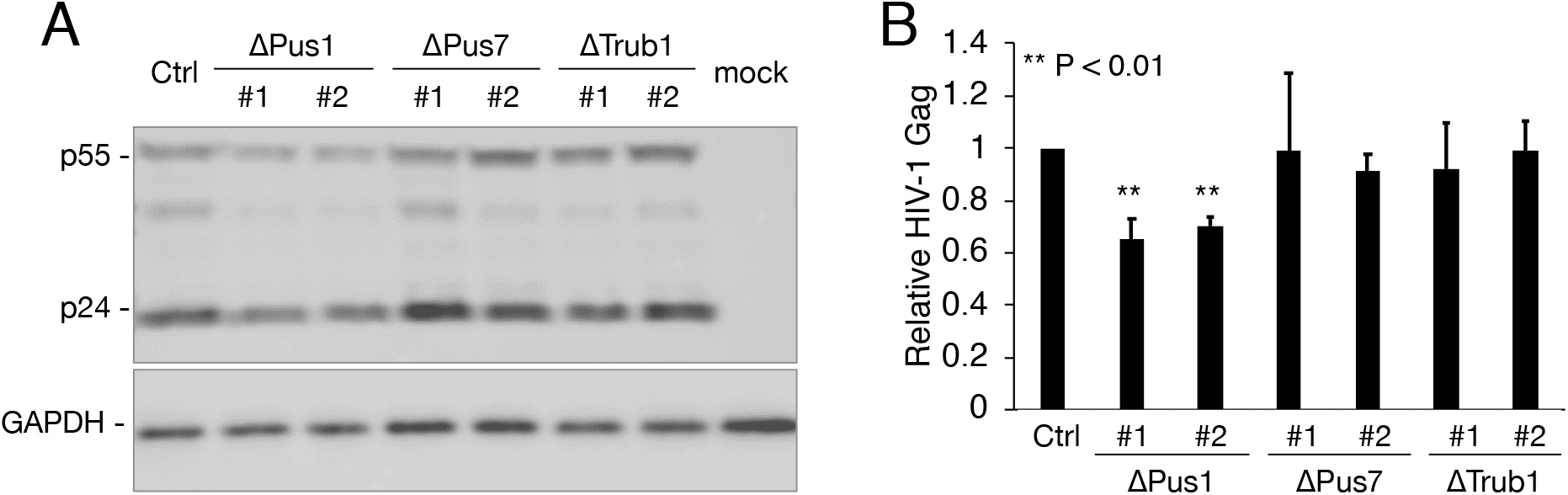
Effect of loss of PUS enzymes on HIV-1 gene expression. A) Western blot of p55 and p24 Gag produced after infection of WT or PUS mutant cell lines at 48 hours post-infection (hpi) with VSV-G pseudotyped NL4-3ΔEnv virions. Cellular GAPDH was used as loading control. B) Similar to panel A except this shows the quantification of total (p55 plus p24) HIV-1 Gag production in HIV-1-infected WT or PUS mutant 293T cell lines at 48hpi across 3 independent biological replicates. Data were normalized to cellular GAPDH and are given relative to the parental 293T cells, which were set at 1.0. ** p<0.01.

## Discussion

The aims of the current study were to 1) develop and validate an antibody based technique that would allow the accurate mapping of Ψ residues on mRNAs with a low input RNA requirement 2) Map any Ψ residues present on HIV-1 transcripts 3) Define the PUS enzyme(s) that deposit these Ψ residues on HIV-1 RNAs by genetic ablation of key human PUS enzymes and 4) Define the phenotypic effect of loss of Ψ residues on HIV-1 RNA function and innate immune activation. We were in fact able to develop an antibody based Ψ mapping technique, called PA-Ψ-seq, and could show that this not only allowed us to confirm the location of known Ψ residues on a range of ncRNAs and mRNAs (Fig.1) but also reproducibly map Ψ residues on HIV-1 transcripts (Fig. 2). We were also able to derive human cell lines in which the three PUS enzymes previously reported (Li et al. 2015; Safra et al. 2017; Borchardt et al. 2020) to add Ψ residues to human mRNAs, PUS1, PUS7 and TRUB1, were individually knocked out using CRISPR/Cas (Fig. 3A-C and Supp. Fig. 2). However, analysis of highly purified mRNA samples isolated from these knock-out clones (Supp. Fig. 4) failed to reveal a significant drop in the Ψ level on mRNA below the ∼0.1% level seen in WT cells (Fig. 3D) and we also did not detect the loss of any of the Ψ residues mapped on HIV-1 RNA (Fig. 2), though a small number of PUS1, PUS7 or TRUB1-dependent sites on cellular mRNAs could be identified (Fig. 4 and Supp. Table 1). How these Ψ are added to HIV-1 transcripts, and whether their addition indeed positively affects HIV-1 gene expression and replication, as we hypothesize, therefore currently remains unclear. However, our data do indicate that the PUS enzymes that add the preponderance, that is more than 50%, of Ψ residues to cellular mRNAs are not PUS1, PUS7 or TRUB1, as previously proposed based largely on *in vitro* analysis or by analogy to yeast PUS enzymes (Carlile et al. 2014; Schwartz et al. 2014; Safra et al. 2017; Martinez et al. 2020), but rather remain to be identified. It also is possible that no single PUS enzyme is responsible for adding the majority of Ψ residues to mRNAs in human cells and that this in fact results from the combined efforts of several of the 12 known human PUS enzymes, including PUS1, PUS7 and TRUB1.

## Materials and Methods

### Cell lines

The Kidney epithelial cell line of human female origin HEK293T (CRL-11268) was cultured in Dulbecco’
ss Modified Eagle Medium (DMEM) supplemented with 7% fetal bovine serum (FBS) and 1% Antibiotic-Antimycotic (Gibco) at 37°C with 5% CO_2_.

### Generation of PUS1, PUS7 and TRUB1 Knock out 293T clones

PUS1, PUS7 and TRUB1 knock out (KO) cell lines were generated by transfecting 293T cells with pLentiCRISPRv2 (Addgene Cat #52961), encoding Cas9 and the following single guide RNAs (sgRNAs). Two PUS1 KO clones (ΔPUS1-1 and ΔPUS1-2) were generated using a single guide RNA (sgRNA) (5’-GAGCCGCATGCCCCCAGGAC-3’) that targets exon 2 in the *PUS1* gene (Li et al. 2015); two PUS7 KO clones (ΔPUS7-1 and ΔPUS7-2) were generated using an sgRNA (5’-TTAATATTGAAACCCCGCTC-3’) obtained from the Gecko Library (Sanjana et al. 2014) that targets exon 10 in the *PUS7* gene; two TRUB1 KO clones (ΔTRUB1-1 and ΔTRUB1-2) were generated using two sgRNAs (5’-CAAAAGTATGGCCGCTTCTG-3’ and 5’-TTCGCCGTGCACAAGCCCAA-3’) obtained from the Gecko Library (Sanjana et al. 2014) to delete a 240 bp region in exon 1 of the *TRUB1* gene (Figure S1). Control cell lines were generated using a non-targeting sgRNA specific for GFP (5’-GTAGGTCAGGGTGGTCACGA-3’). The polyclonal cultures were then single cell cloned, assessed for target protein expression by Western blot. To validate CRISPR mutations, the genomic DNA from all KO cell lines was extracted and the region flanking the Cas9 target site from each ΔPUS cell line PCR amplified and cloned into the EcoRI and HindIII sites of pGEM-3zf vector (Promega). The primers used to amplify the region flanking the Cas9 target sites were the following: PUS1 forward: 5’-GTCAGGGGTCAGAAGGAACAG-3’ and reverse: 5’-TCACCCTGATCACCGCAAAC-3’; PUS7 forward: 5’-ATGAGATGATGTAGGACCAGGTG-3’ and reverse: 5’-TCCTCAAGGTGTTTTTGCCAAGT-3’; TRUB1 forward: 5’-AAGGCCATGGACTACAATTC-3’ and reverse: 5’-CCTGGAGCAACTATTTAGAATT-3’. At least 10 independent clones were isolated and used for Sanger sequencing.

### HIV-1 Infections

Wild type HIV-1 isolate NL4-3 was generated by transfection of 293T cells with pNL4-3 (NIH AIDS Reagent Program, Cat#114) using polyethylenimine (PEI). The media were replaced the next day, supernatant media collected at 72h post transfection (hpi) and filtered through a 0.45µm filter before being used to infect cells. The pNL4-3ΔEnv plasmid expresses an NL4-3 provirus that is full length except that it bears a 949 base pair deletion of the Env ORF that also introduces a frameshift mutation that inactivates the remaining ORF. This deletion removes Env amino acids (aa) 29 to 345, leaving only a vestigial 28aa product. pNL4-3ΔEnv DNA was co-transfected into 293T cells using PEI together with pMD2.G, which encodes the VSV-G glycoprotein. 72 hpi the virus containing supernatant medium was collected and passed through a 0.45 µM filter before use in subsequent infection experiments. 500,000 293T cells (WT, ΔPUS1-1, ΔPUS1-2, ΔPUS7-1, ΔPUS7-2, ΔTRUB1-1, and ΔTRUB1-2) were resuspended in 1.5 ml of media either lacking or containing VSV-G pseudotyped pNL4-3ΔEnv virus. The infected cells were collected at 48 hpi and analysed by Western blot.

### Western blots

Protein samples were collected at 48 hpi and lysed using Laemmli buffer. The samples were sonicated, denatured at 95°C for 10 minutes and separated on Tris-Glycine-SDS polyacrylamide gels. Proteins were transferred to a nitrocellulose membrane, blocked with 5% milk in PBS with 0.1% Tween (PBST) and then incubated with primary and secondary antibodies diluted in PBST. Western blots used the following antibodies HIV-1 p24 (AIDS Reagent Program-6458), PUS1 (Proteintech #11512-1-AP), PUS7 (Abcam, #ab226257), TRUB1 (Aviva Systems Biology #ARP62931_P050) and GAPDH (Proteintech #60004-1.1g) loading control.

### PA-Ψ-seq

PA-Ψ-seq was performed as described previously (Chen et al. 2015; Courtney et al. 2019c; Tsai et al. 2020) with some modifications. The 293T, ΔPUS1, ΔPUS7 or ΔTRUB1 cell lines were transfected with 10µg of a plasmid encoding CD4 (pcDNA-CD4). Next day, the media was replaced. Cells were split and infected with NL4-3 virus for 48h and then pulsed with 100µM of either 4SU (Carbosynth Cat #NT06186) or 6SG (Sigma Cat #858412) for a further 24h. Total poly(A) RNA was purified using magnetic beads (Poly(A)Purist MAG Kit, Invitrogen #AM1922) following the manufacturer’s instructions. The immunoprecipitation (IP) was performed using an anti-pseudouridine (anti-Ψ) antibody (MBL #347-3) while the m^6^A control was performed using an anti-m^6^A antibody (Synaptic Systems #202-003). 10µg of Poly(A) RNA was incubated with 5µg of the RNA modification specific antibody and RNasin (40U/µl) for 2h, crosslinked with UV 365nm 2500 x100µJ/cm^2^ twice in a UV Stratalinker, and then treated with RNase T1 (0.1U/µl) for 15min at 22°C. Poly(A) RNA-antibody complexes were immunoprecipitated using Protein G magnetic beads (Thermo Dynabeads #10004D) for 1h at 4°C in rotation. Next, RNA-antibody-protein G bead complexes were washed with IP buffer (3X) and incubated with RNase T1 (15U/µl) for a further 15min at 22°C. RNA end repair was performed by incubating the poly (A) RNA-antibody-protein G beads with CIP (0.5 U/µl) for 10min at 37°C. After successive washes with phosphatase wash buffer (2X), and PNK buffer (2X), RNA was phosphorylated by adding T4-PNK (1U/µl) in PNK buffer with DTT (5mM) and ATP (10mM) for 30min at 37°C. The complexes were washed with PNK buffer (3X) and RNA was eluted from the beads by incubation with Proteinase K, 90min at 55°C. RNA was extracted using Trizol (Invitrogen). The RNA recovered from the IP was used for cDNA library preparation using the NEB Next Small RNA Library Prep Set for Illumina (NEB, E7330S) according to the manufacturer instructions.

For virion RNA, HIV-1 virions were isolated as previously described (Eckwahl et al. 2016; Courtney et al. 2019c; Tsai et al. 2020). The supernatant from HIV-1 infected, 4SU/6SG pulsed 293T, ΔPUS1, ΔPUS7 or ΔTRUB1 cells was collected at 72 hpi, filtered through a 0.45µm filter and pelleted by ultracentrifugation (38,000 rpm for 90 min) through a 20% sucrose cushion. Total RNA was then extracted using TRIzol and the IP was performed as described above.

### Bioinformatics

High throughput sequencing data were analyzed as previously reported (Tsai et al. 2020). Fastx toolkit (Gordon and Hannon 2010) was used to remove adaptor sequences while reads shorter than 15nt were discarded. Pass filter reads were then aligned to Human genome build hg19 using Bowtie (Langmead et al. 2009), allowing 1 mismatch. Unaligned reads were then aligned to HIV-1 NL4-3 with a single copy of the long terminal repeat (U5 on the 5’ end, and U3-R on the 3’ end, essentially 551-9626 nt of GenBank AF324493.2.), also allowing 1 mismatch. In-house Perl scripts were used to only retain aligned reads that contain the characteristic T>C or G>A mutations resulting from 4SU or 6SG incorporation. Finally, Samtools (Li et al. 2009) was used to convert alignment file formats to be suitable for visualization with IGV (Robinson et al. 2011).

PA-Ψ-seq was validated by aligning sequencing reads to human ribosomal 28S, 18S and 5.8S rRNAs and compared to previously reported Ψ sites (Lestrade and Weber 2006; Taoka et al. 2018). The sequences and the coordinates of previously reported Ψ sites in mRNA on the PIANO Database (Song et al. 2020) were compared to the Ψ sites present in our datasets. RNA reads were also aligned to tRNA and snRNA sequences (Lowe and Eddy 1997) and compared to previously reported Ψ sites (Jühling et al. 2009; Bohnsack and Sloan 2018). tRNA sequences used for alignment were from tRNAdb (Jühling et al. 2009), the rRNA sequence used was the same as used in snoRNABase (Lestrade and Weber 2006): 28S rRNA: GenBank #U13369 nts 7935-12969, 18S rRNA: X03205, 5.8S rRNA: U13369 nts 6623-6779. The snRNA sequences were downloaded individually from GenBank human genome build GRCh38.p13. Peak calling on cellular RNAs was performed using MACS2 (Zhang et al. 2008). The distribution of the Ψ sites on transcripts was analyzed using MetaplotR (Olarerin-George and Jaffrey 2017).

For *de novo* identification of PUS1/PUS7/TRUB1 dependent Ψ sites, sequencing reads from PA-Ψ-seq runs of cellular poly(A) RNA+ were aligned to the hg19 human transcriptome using Tophat (Trapnell et al. 2009) (transcript annotation file downloaded from the UCSC genome browser), allowing one mismatch. Discovery of Pus-dependent sites was done by subtracting the Ψ sites present in the KO data set from a paired WT control data set. To minimize the possibility of wrongly calling Ψ sites, where the local read depth was diminished but not gone in the Pus KO data set, as Pus-dependent, we took a conservative approach, using a higher peak calling threshold for the WT data sets than the KO data sets. To ensure a comparison of equal read depth between WT and KO data sets, the read depths of WT-KO pairs (IP’ed side by side) were normalized by downsampling the one with the higher total read depth to the same as the one with a lower read depth using the “randsample” function of MACS2. All peak calling was done using MACS2 allowing a Q value (minimum false discovery rate cutoff) of 0.0005 (macs2 callpeak--nomodel -q 0.0005 --extsize 32 --shift 0 --keep-dup 5 -g hs), while allowing a Q value of 0.05 for all KO data sets. The MACS2 called site lists were then passed through a filter while converting the xls file to a bam file (awk ‘{if($1!=“#” && $1!=“chr” && $6>=3 && $8>=5) {print $1 “\t” $2-1 “\t” $3 “\t” $10 “\t” $6 “\t.”}}’ peaks.xls > filtered_peaks.bed), only allowing sites with at least 3x pileup and 5x fold enrichment and trashing all other sites. For KO data sets, this filter criteria was loosened to allow sites with more than 1x pileup read count and 3x fold enrichment. Using the intersect function of Bedtools (Quinlan 2014), we first looked for Ψ sites consistent across an earlier independent run and the KO-paired WT run, to only consider repeatable sites. These repeatable sites were then again screened for KO-absent sites using Bedtools intersect. All Ψ site overlaps defined as at least 33% overlap. Lastly, the PUS/TRUB-dependent site list was manually curated by the following criteria: PUS7 dependent sites were listed only if there were 0 reads in the KO. PUS1/TRUB1 dependent sites were listed only if the read depth normalized read count KO/WT ratio of both repeats (4SU&6SG) was < 0.2

### RNA Modification Quantification by UPLC-MS/MS

PolyA+ mRNA was isolated from total RNA using two rounds of oligo-dT affinity purification (NEBNext Poly(A) mRNA Magnetic Isolation Module), followed by RiboZero treatment (Illumina). RNA quality was assessed using a Bioanalyzer 2100 with RNA 6000 Nano kit (Agilent). 500ng of total RNA or mRNA for each sample were then digested to single nucleosides as previously described (Courtney et al. 2019b). Briefly, RNA was incubated with nuclease P1 (Sigma, 2U) in buffer containing 25mM NaCl and 2.5mM ZnCl2 for 2h at 37°C, followed by incubation with Antarctic Phosphatase (NEB, 5U) for an additional 2h at 37°C. UPLC-MS/MS quantification of modified and unmodified nucleosides was modified from a previously published protocol (Basanta-Sanchez et al. 2016). Stable isotope-labeled internal standards ^13^C_5_-adenosine (Cambridge Isotope Labs, CLM-3678-PK) and ^15^N_3_-cytidine (Cambridge Isotope Labs, NLM-3797-PK) were spiked into each sample at a final concentration of 100nM. UPLC-MS/MS was performed using a Waters Acquity UPLC and Xevo TQ-XS mass spectrometer in MRM mode, with the following details: 2.1 x 50 mm HSS T3 C18 1.7 um column, 0.01% formic acid in water as mobile phase A with a gradient to 50/50 MeCN/Water containing 0.01% formic acid with mobile phase B. Calibration curves for modified and unmodified nucleoside standards were linear over five orders of magnitude.

## Data availability

Deep sequencing data from the PA-Ψ-seq assays have been deposited at the NCBI GEO database under accession number GSE172136.

## Acknowledgements

This research was funded in part by NIH grant R01-DA046111 to B.R.C. and a Duke University Center for AIDS Research (CFAR, P30-AI064518) pilot award to K.T. B.R.C was supported by the Center for HIV RNA Studies (CRNA, NIH award U54-AI15047). The following reagents were obtained through the NIH AIDS Reagent Program, Division of AIDS, NIAID, NIH: HIV-1 p24 Gag Monoclonal (#24-3) from Michael Malim. C.M.C. was funded in part by the Mexican National Science and Technology Council (CONACyT) fellowship: 711023/740541. We thank the Duke University School of Medicine for the use of the Proteomics and Metabolomics Shared Resource, which provided UPLC-MS/MS services.

## Supplementary Figure Legends

**Supplementary Figure 1.**
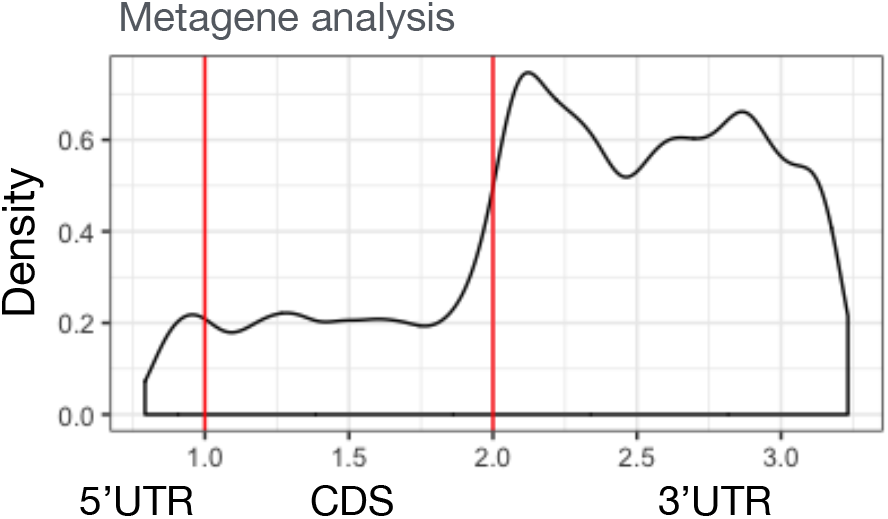
This figure shows a metagene analysis of the relative location of Ψ residues across the 5’UTR, CDS and 3’UTR of cellular mRNA species. Peaks from PA-Ψ-seq of 293T cell poly(A)+ mRNAs were called using MACS2, and the called peak distribution on cellular transcripts analyzed by MetaplotR.

**Supplementary Figure 2.**
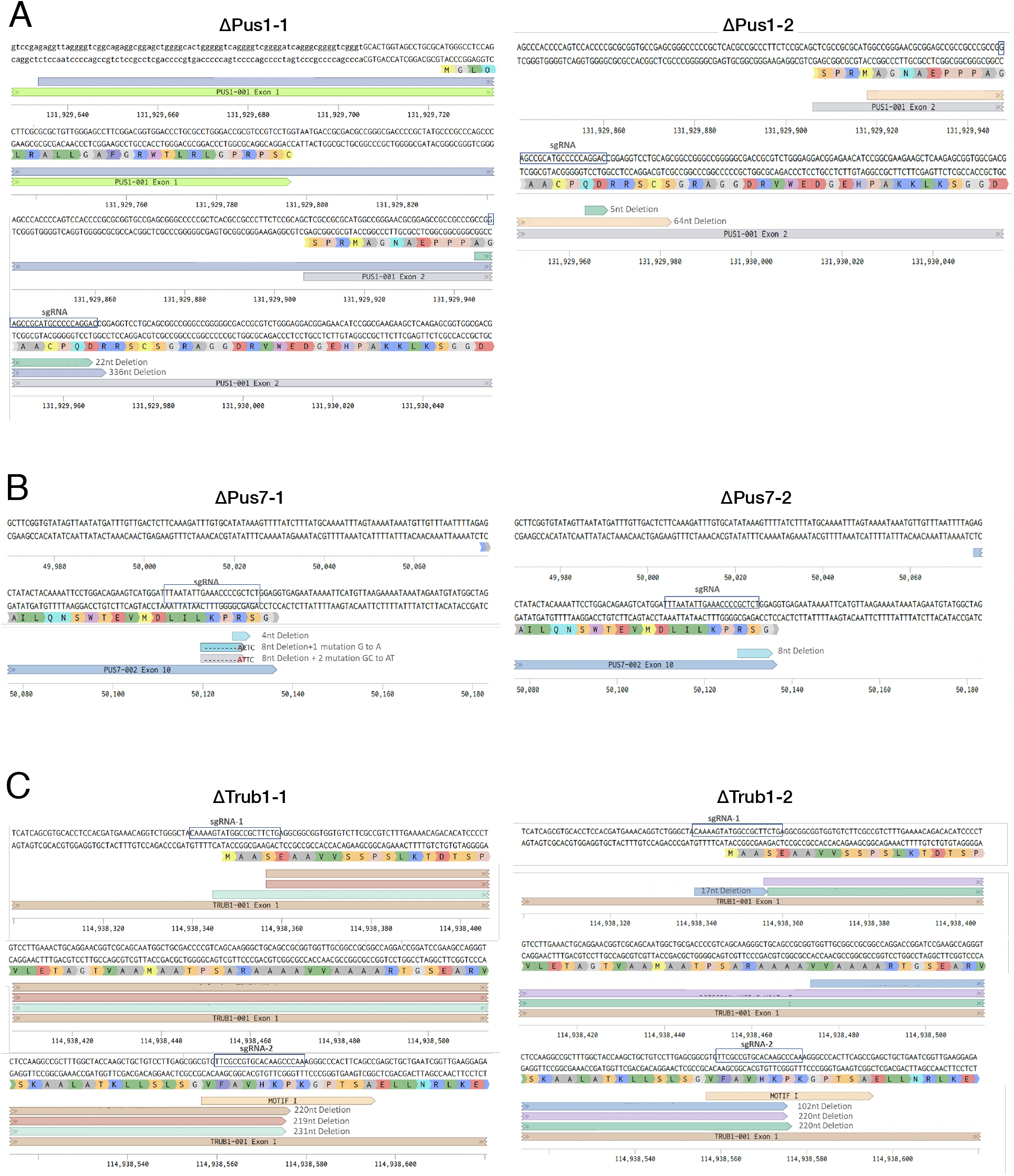
This figure presents the DNA sequence analysis of the mutations introduced into the genomic copies of the *pus1, pus7* and *trub1* gene in 293T cells by gene editing using CRISPR/Cas. The location of relevant coding exons and the location and extent of introduced deletion and missense mutations is indicated.

**Supplementary Figure 3.**
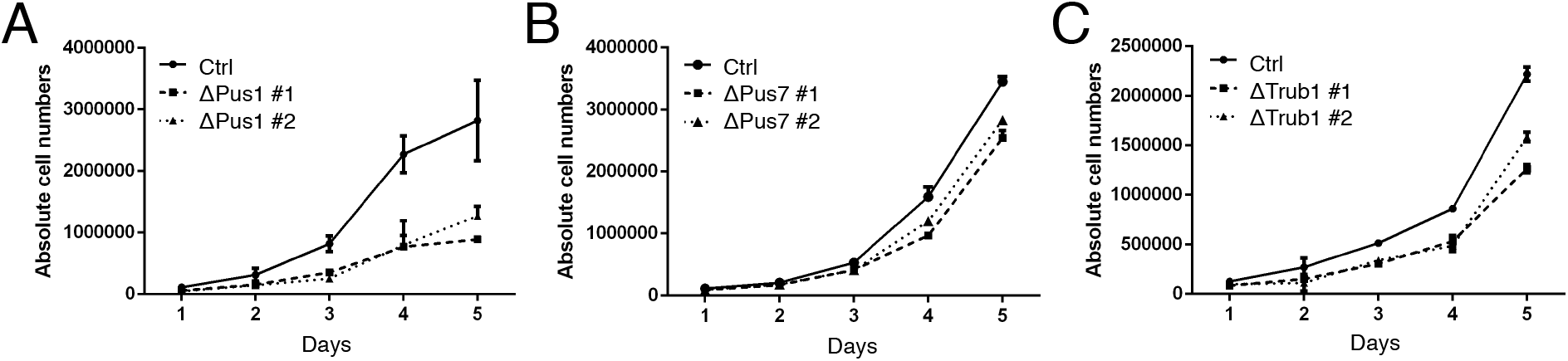
Growth curves comparing the rate of growth of the indicated mutant 293T clonal cell lines with wildtype 293T cells. Live cells were counted daily over a 5-day period. Average of three biological replicates with SD indicated.

**Supplementary Figure 4.**
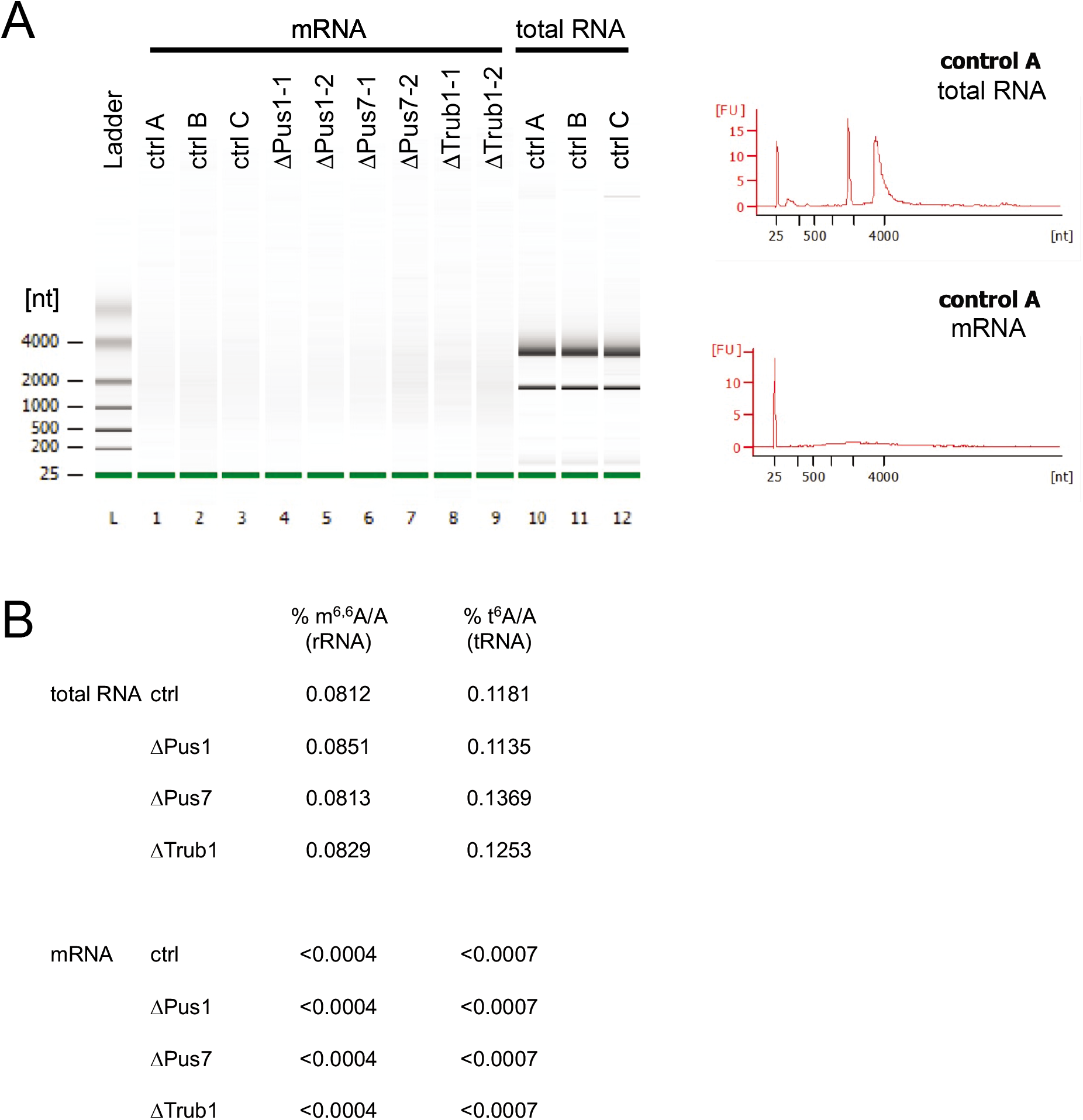
Validation of mRNA sample purity. Total RNA was subjected to two rounds of oligo-dT affinity purification to capture polyA mRNA, followed by RiboZero rRNA depletion. (A) Bioanalyzer 2100 analysis of RNA samples, including representative electropherograms. L = ladder, lanes 1-9 show mRNA samples as annotated in image, lanes 10-12 show total RNA prior to mRNA isolation. Note the absence of rRNA bands or peaks in highly purified mRNA. (B) Modified nucleosides were measured from total and mRNA samples by UPLC-MS/MS. Average m^6,6^A and t^6^A levels are shown as % of the unmodified nucleotide (mod/unmod*100); these modifications are specific for rRNA and tRNA, respectively. Neither modification was detected above the lower limit of detection in the highly purified mRNA samples.

**Supplementary Table 1:** List of identified PUS1, PUS7 or TRUB1-dependent sites of Ψ addition on cellular mRNAs.

## References

Anderson BR, Muramatsu H, Nallagatla SR, Bevilacqua PC, Sansing LH, Weissman D, Karikó K. 2010. Incorporation of pseudouridine into mRNA enhances translation by diminishing PKR activation. Nucleic acids research 38: 5884–5892.

Basanta-Sanchez M, Temple S, Ansari SA, D’Amico A, Agris PF. 2016. Attomole quantification and global profile of RNA modifications: Epitranscriptome of human neural stem cells. Nucleic acids research 44: e26.

Bohnsack MT, Sloan KE. 2018. Modifications in small nuclear RNAs and their roles in spliceosome assembly and function. Biological chemistry 399: 1265–1276.

Borchardt EK, Martinez NM, Gilbert WV. 2020. Regulation and Function of RNA Pseudouridylation in Human Cells. Annual review of genetics 54: 309–336.

Carlile TM, Rojas-Duran MF, Gilbert WV. 2015. Pseudo-Seq: Genome-Wide Detection of Pseudouridine Modifications in RNA. Methods in enzymology 560: 219–245.

Carlile TM, Rojas-Duran MF, Zinshteyn B, Shin H, Bartoli KM, Gilbert WV. 2014. Pseudouridine profiling reveals regulated mRNA pseudouridylation in yeast and human cells. Nature 515: 143–146.

Chen K, Lu Z, Wang X, Fu Y, Luo GZ, Liu N, Han D, Dominissini D, Dai Q, Pan T et al. 2015. High-resolution N(6) -methyladenosine (m(6) A) map using photo-crosslinking-assisted m(6) A sequencing. Angewandte Chemie 54: 1587–1590.

Courtney DG, Chalem A, Bogerd HP, Law BA, Kennedy EM, Holley CL, Cullen BR. 2019a. Extensive Epitranscriptomic Methylation of A and C Residues on Murine Leukemia Virus Transcripts Enhances Viral Gene Expression. mBio 10.

Courtney DG, Tsai K, Bogerd HP, Kennedy EM, Law BA, Emery A, Swanstrom R, Holley CL, Cullen BR. 2019b. Epitranscriptomic Addition of m5C to HIV-1 Transcripts Regulates Viral Gene Expression. Cell host & microbe 26: 217-227.e216.

Courtney DG, Tsai K, Bogerd HP, Kennedy EM, Law BA, Emery A, Swanstrom R, Holley CL, Cullen BR. 2019c. Epitranscriptomic Addition of m(5)C to HIV-1 Transcripts Regulates Viral Gene Expression. Cell host & microbe 26: 217–227 e216.

Durbin AF, Wang C, Marcotrigiano J, Gehrke L. 2016. RNAs Containing Modified Nucleotides Fail To Trigger RIG-I Conformational Changes for Innate Immune Signaling. mBio 7.

Eckwahl MJ, Arnion H, Kharytonchyk S, Zang T, Bieniasz PD, Telesnitsky A, Wolin SL. 2016. Analysis of the human immunodeficiency virus-1 RNA packageome. Rna 22: 1228–1238.

Gilbert WV, Bell TA, Schaening C. 2016. Messenger RNA modifications: Form, distribution, and function. Science (New York, NY) 352: 1408–1412.

Gordon A, Hannon G. 2010. FastX toolkit.

Hafner M, Landthaler M, Burger L, Khorshid M, Hausser J, Berninger P, Rothballer A, Ascano M, Jr., Jungkamp AC, Munschauer M et al. 2010. Transcriptome-wide identification of RNA-binding protein and microRNA target sites by PAR-CLIP. Cell 141: 129–141.

Hughes DG, Maden BE. 1978. The pseudouridine contents of the ribosomal ribonucleic acids of three vertebrate species. Numerical correspondence between pseudouridine residues and 2’-O-methyl groups is not always conserved. The Biochemical journal 171: 781–786.

Jühling F, Mörl M, Hartmann RK, Sprinzl M, Stadler PF, Pütz J. 2009. tRNAdb 2009: compilation of tRNA sequences and tRNA genes. Nucleic acids research 37: D159–162.

Karijolich J, Yu YT. 2011. Converting nonsense codons into sense codons by targeted pseudouridylation. Nature 474: 395–398.

Kariko K, Buckstein M, Ni H, Weissman D. 2005. Suppression of RNA recognition by Toll-like receptors: the impact of nucleoside modification and the evolutionary origin of RNA. Immunity 23: 165–175.

Karikó K, Muramatsu H, Welsh FA, Ludwig J, Kato H, Akira S, Weissman D. 2008. Incorporation of pseudouridine into mRNA yields superior nonimmunogenic vector with increased translational capacity and biological stability. Molecular therapy : the journal of the American Society of Gene Therapy 16: 1833–1840.

Kennedy EM, Bogerd HP, Kornepati AV, Kang D, Ghoshal D, Marshall JB, Poling BC, Tsai K, Gokhale NS, Horner SM et al. 2016. Posttranscriptional m(6)A Editing of HIV-1 mRNAs Enhances Viral Gene Expression. Cell host & microbe 19: 675–685.

Langmead B, Trapnell C, Pop M, Salzberg SL. 2009. Ultrafast and memory-efficient alignment of short DNA sequences to the human genome. Genome biology 10: R25.

Lestrade L, Weber MJ. 2006. snoRNA-LBME-db, a comprehensive database of human H/ACA and C/D box snoRNAs. Nucleic acids research 34: D158–162.

Li H, Handsaker B, Wysoker A, Fennell T, Ruan J, Homer N, Marth G, Abecasis G, Durbin R, Genome Project Data Processing S. 2009. The Sequence Alignment/Map format and SAMtools. Bioinformatics 25: 2078–2079.

Li X, Zhu P, Ma S, Song J, Bai J, Sun F, Yi C. 2015. Chemical pulldown reveals dynamic pseudouridylation of the mammalian transcriptome. Nature chemical biology 11: 592–597.

Lowe TM, Eddy SR. 1997. tRNAscan-SE: a program for improved detection of transfer RNA genes in genomic sequence. Nucleic acids research 25: 955–964.

Martinez NM, Su A, Nussbacher JK, Burns MC, Schaening C, Sathe S, Yeo GW, Gilbert WV. 2020. Pseudouridine synthases modify human pre-mRNA co-transcriptionally and affect splicing. bioRxiv: 2020.2008.2029.273565.

Meyer KD, Jaffrey SR. 2014. The dynamic epitranscriptome: N6-methyladenosine and gene expression control. Nature reviews Molecular cell biology 15: 313–326.

Nance KD, Meier JL. 2021. Modifications in an Emergency: The Role of N1-Methylpseudouridine in COVID-19 Vaccines. ACS Central Science.

Olarerin-George AO, Jaffrey SR. 2017. MetaPlotR: a Perl/R pipeline for plotting metagenes of nucleotide modifications and other transcriptomic sites. Bioinformatics 33: 1563–1564.

Quinlan AR. 2014. BEDTools: The Swiss-Army Tool for Genome Feature Analysis. Current protocols in bioinformatics 47: 11 12 11–34.

Rintala-Dempsey AC, Kothe U. 2017. Eukaryotic stand-alone pseudouridine synthases - RNA modifying enzymes and emerging regulators of gene expression? RNA biology 14: 1185–1196.

Robinson JT, Thorvaldsdottir H, Winckler W, Guttman M, Lander ES, Getz G, Mesirov JP. 2011. Integrative genomics viewer. Nat Biotechnol 29: 24–26.

Roundtree IA, Evans ME, Pan T, He C. 2017. Dynamic RNA Modifications in Gene Expression Regulation. Cell 169: 1187–1200.

Safra M, Nir R, Farouq D, Vainberg Slutskin I, Schwartz S. 2017. TRUB1 is the predominant pseudouridine synthase acting on mammalian mRNA via a predictable and conserved code. Genome research 27: 393–406.

Sanjana NE, Shalem O, Zhang F. 2014. Improved vectors and genome-wide libraries for CRISPR screening. Nat Methods 11: 783–784.

Schwartz S, Bernstein DA, Mumbach MR, Jovanovic M, Herbst RH, León-Ricardo BX, Engreitz JM, Guttman M, Satija R, Lander ES et al. 2014. Transcriptome-wide mapping reveals widespread dynamic-regulated pseudouridylation of ncRNA and mRNA. Cell 159: 148–162.

Song B, Tang Y, Wei Z, Liu G, Su J, Meng J, Chen K. 2020. PIANO: A Web Server for Pseudouridine-Site (Ψ) Identification and Functional Annotation. Frontiers in genetics 11: 88.

Taoka M, Nobe Y, Yamaki Y, Sato K, Ishikawa H, Izumikawa K, Yamauchi Y, Hirota K, Nakayama H, Takahashi N et al. 2018. Landscape of the complete RNA chemical modifications in the human 80S ribosome. Nucleic acids research 46: 9289–9298.

Trapnell C, Pachter L, Salzberg SL. 2009. TopHat: discovering splice junctions with RNA-Seq. Bioinformatics 25: 1105–1111.

Tsai K, Cullen BR. 2020. Epigenetic and epitranscriptomic regulation of viral replication. Nat Rev Microbiol 18: 559–570.

Tsai K, Jaguva Vasudevan AA, Martinez Campos C, Emery A, Swanstrom R, Cullen BR. 2020. Acetylation of Cytidine Residues Boosts HIV-1 Gene Expression by Increasing Viral RNA Stability. Cell host & microbe 28: 306–312 e306.

Wiener D, Schwartz S. 2021. The epitranscriptome beyond m(6)A. Nature reviews Genetics 22: 119–131.

Yue Y, Liu J, He C. 2015. RNA N6-methyladenosine methylation in post-transcriptional gene expression regulation. Genes & development 29: 1343–1355.

Zhang W, Eckwahl MJ, Zhou KI, Pan T. 2019. Sensitive and quantitative probing of pseudouridine modification in mRNA and long noncoding RNA. RNA (New York, NY) 25: 1218–1225.

Zhang Y, Liu T, Meyer CA, Eeckhoute J, Johnson DS, Bernstein BE, Nusbaum C, Myers RM, Brown M, Li W et al. 2008. Model-based analysis of ChIP-Seq (MACS). Genome biology 9: R137.

Zhou J, Liang B, Li H. 2010. Functional and structural impact of target uridine substitutions on the H/ACA ribonucleoprotein particle pseudouridine synthase. Biochemistry 49: 6276–6281.

Zucchini C, Strippoli P, Biolchi A, Solmi R, Lenzi L, D’Addabbo P, Carinci P, Valvassori L. 2003. The human TruB family of pseudouridine synthase genes, including the Dyskeratosis Congenita 1 gene and the novel member TRUB1. Int J Mol Med 11: 697–704.

